# Complex genetic determinism of male-fertility restoration in the gynodioecious snail *Physa acuta*

**DOI:** 10.1101/2024.08.08.607164

**Authors:** Elpida Skarlou, Fanny Laugier, Kévin Béthune, Timothée Chenin, Jean-Marc Donnay, Céline Froissard, Patrice David

## Abstract

Male fertility in plants is often controlled by the interaction between mitochondrial and nuclear genes. Some mitotypes confer cytoplasmic male sterility (CMS), making the individual male-sterile, unless the nuclear background contains alleles called restorers, that suppress the effects of CMS and restore the hermaphroditic phenotype. Restorers in cultivated crops are often alleles with strong and dominant effect, but in wild plants, data often suggest more complex systems. Here, we characterized the inheritance and specificity of restoration in a new CMS model, the freshwater snail *Physa acuta*. We explored two different populations (i) a naive population i.e., without contact with CMS in the past 80 generations, (ii) a non-naive population, where CMS is present and largely restored. Although we found male fertility of individuals with CMS mitogenomes to be heritable in both contexts, this genetic determinism was of a different nature depending on population history. In naive populations not coevolved with CMS the background variation may include alleles that happen to act as weak quantitative modifiers of the penetrance of CMS, while in populations coevolved with CMS, selection may have favored, when such variants were available, the emergence of strong alleles with a dominant effect.

## Introduction

Gynodioecy is a sexual polymorphism that is common in angiosperms (Dufay et al., 2014; Caruso et al., 2016), in which hermaphrodites coexist with male-sterile individuals (i.e., functionally females, Saumitou-Laprade et al., 1994). Gynodioecy represents a well-known example of genetic conflict (Burt & Trivers, 2006) because sex determinism in gynodioecious systems is often cyto-nuclear with cytoplasmic male sterility (CMS) genes, associated with the mitochondrial genome, suppressing pollen production and rendering an individual functionally female unless one or more nuclear genes restore the male function (Cosmides & Tooby, 1981; Saumitou-Laprade et al., 1994; Werren & Beukeboom, 1998). An individual is phenotypically female if it carries CMS without one or more CMS-specific nuclear genes (i.e., nuclear restorers). It is a hermaphrodite if it carries either (i) a male-fertile cytoplasm, or (ii) a CMS cytoplasm and a suitable nuclear restorer. CMS has been documented in 140 species of angiosperms (Laser & Lersten, 1972), and 28 CMS genes from 13 crop species have been identified (Chen & Liu, 2014).

It appears that cyto-nuclear gynodioecious systems often contain multiple forms of CMS (e.g. *Oryza sativa, Zea mays*, see Chen and Liu 2014 for a review), each requiring its own mode of restoration (de Haan et al., 1997; Charlesworth & Laporte, 1998; van Damme et al., 2004). Fourteen restorer genes (*Rf*) from nine crops have been isolated (see Table 1 Kim & Zhang, 2018), with the majority of them encoding PPR proteins (Pentatricopeptide Protein Repeat). However, *Rf* genes encoding other proteins have been discovered, including peptidase in sugar beet (Yamamoto et al., 2008; Matsuhira et al., 2012) and aldehyde dehydrogenase in maize (Cui et al., 1996). In most cases, a *Rf* gene is specific to a single CMS, whereas a CMS gene can be affected by multiple *Rf* genes (see Table 1 Chen and Liu 2014). For example, in *Z. mays* the CMS-T is restored by both *Rf1* and *Rf2* (Cui et al., 1996; Dewey et al., 1987; Dill et al., 1997) and in *Plantago coronopus* at least five restorer alleles seem to be involved in the restoration of CMS (Koelewijn & Van Damme, 1995). Cases in which a gene restores more than one CMS mitotype are rare. Such an example is found in the common rice, where *Rf5* and *Rf6* (initially identified in CMS-HL individuals) are also able to restore CMS-BT mitotype (Huang et al., 2012, 2015; Kim & Zhang, 2018).

**Table 1:** Sample sizes by category. For each combination of paternal and maternal types, we provide N_1_/N_2_, where N_1_ is the number of G_1_ individuals successfully assessed for male fertility, and N_2_ is the number of G_0_ mothers (each mother produced 1 to 3 G_1_ individuals).

|  |  | Maternal |  |  |  |
| --- | --- | --- | --- | --- | --- |
|  |  | KS | KF | DS | Total |
| Paternal | Lyon inbred lines | 164/82 | 36/21 | 65/47 | 265/150 |
|  | Montpellier inbred lines | 350/231 | 220/130 | 211/168 | 781/529 |
|  | Total inbred lines | 524/313 | 256/140 | 276/215 | 1056/668 |
|  | HFR selected population | 52/31 | 49/27 | 32/22 | 133/80 |
|  | LFR selected population | 248/105 | 39/20 | 32/25 | 319/150 |
|  | Total selected populations | 300/136 | 88/47 | 64/47 | 452/230 |

The maintenance of cyto-nuclear polymorphism has been extensively modelled (e.g. (Charlesworth & Ganders, 1979; Couvet et al., 1998; Gouyon et al., 1991; Bailey et al., 2003; Dufaÿ et al., 2007) and theoretical studies have shown that the stability of this polymorphism depends on a variety of factors, including the positive pleiotropic effects of CMS alleles (i.e., seed fitness advantage of females, or “female advantage”, Lewis, 1941) and negative pleiotropic effects of restorer alleles on either seed and/or pollen fitness (“cost of restoration”, Charlesworth & Ganders, 1979; Delannay et al., 1981; Frank, 1989; Gouyon et al., 1991). Models often assume that restoration of male fertility is achieved through one allele (but see Frank 1989), with variable assumptions regarding its dominance (Delph et al., 2007). Most models considered dominant restorer alleles, with only a couple of cases considering recessive inheritance (Ross & Gregorius, 1985), a modeling approach supported by empirical studies, as most of the crop species exhibited dominant restorer alleles (see Table 2 Delph 2007 for a review but see also *Plantago lanceolata*, van Damme, 1984; de Haan et al., 1997). However, in wild populations, the restoration of male fertility is often more complex, involving several nuclear loci (Charlesworth & Laporte, 1998; Koelewijn, 2003; Touzet et al., 2004, Dufay et al., 2008). For example, Koelewijn & van Damme (1995) suggested that at least five restorer loci were involved in restoring male sterile cytoplasms of *Plantago coronopus.* In addition, Dudle et al. (2001) found that sex determinism in *Lobelia siphilitica* seemed to vary among cytoplasms with one CMS gene being restored by a single dominant allele, while restoration of other CMS could only be explained by the action of several nuclear loci and/or epistatic effects. Inspired by those examples, Ehlers et. al (2005) proposed the use of the quantitative-genetic threshold model to describe sex-determinism in some gynodioecious species. In this model, individuals with a “maleness” trait above a given threshold develop a hermaphroditic phenotype, while those below the threshold develop a female phenotype.

**Table 2:** Generalised linear mixed models on the binomial male-fertility variable (fertile versus sterile or semi-sterile) in the offspring generation, selected lines dataset. LRT tests are given for fixed and random factors (χ²_df_ and corresponding *p-*values). For random factors we also provide estimates of variance (v, on logit scale) and *p*-values account for the one-sided alternative (variances cannot be negative). Note that the very high « maternal ID » variance in the KF category is an effect of the logit scale when proportions are close to 0 or 100% (here 92% individuals are fertile): in these conditions a very large variance in the logit scale makes a small variance in actual proportions. The block random factor was omitted in this table because in all models the estimated variance was 0.00. *p-*values are bolded when <0.05. Note that full-sibs (offspring sharing the same mother and father) tend to have similar phenotypes as attested by significant effects of mother identity on top of what is already explained by the fixed effect of maternal type. This mother effect is not predicted by maternal phenotype (among KS mothers) as it remains approximately the same in the model with the maternal phenotype included as a fixed effect (χ²_1df_ = 4.80, variance = 0.72, *p* = 0.014). The maternal ID effect is also, surprisingly, significant within the KF category. This is because, among the few cases of male sterility observed in individuals with a KF mother and an HFR father (four in total), two share the same mother—a pattern unlikely to have occurred by chance.

| Data | Fixed effects |  |  | random effects |
| --- | --- | --- | --- | --- |
|  | maternal type<br>(DS, KF, KS) | paternal type<br>(HFR, LFR) | Interaction | maternal ID |
| All | $\chi^2_{df2} = 64.36$<br>$p = \mathbf{1.06 \cdot 10^{-14}}$ | $\chi^2_{df1} = 4.25$<br>$p = \mathbf{0.039}$ | $\chi^2_{df2} = 0.91$<br>$p = 0.634$ | $\chi^2_{df1} = 6.79$, $v = 1.01$<br>$p = \mathbf{0.0045}$ |
| only DS | | $\chi^2_{df1} = 0.356$<br>$p = 0.551$ | | $\chi^2_{df1} = 0.00$, $v = 0.00$<br>$p = 0.500$ |
| only KF | | $\chi^2_{df1} = 0.001$<br>$p = 0.970$ | | $\chi^2_{df1} = 10.07$, $v = 91.72$<br>$p = \mathbf{0.008}$ |
| only KS | | $\chi^2_{df1} = 4.23$,<br>$p = \mathbf{0.039}$ | | $\chi^2_{df1} = 6.79$, $v = 0.76$<br>$p = \mathbf{0.010}$ |

A mixture of several factors is also possible, and strong, dominant specific restorers acting on one or a few CMS mitotypes, as found in present-day plant populations, may result from a history of coevolutionary arms-races with their specific CMS genes (e.g. Sloan et al., 2012), and selective sweeps of favorable mutants (Bergero et al., 2019). When a new CMS type first appears, it is unlikely that such strong-effect specific restorer alleles are already present in the population. However, the standing variation may contain genotypes that happen to modulate the penetrance of CMS, not necessarily fully restoring male-fertility, and not necessarily in a specific way. This background variation may often be present, but its effects are not easy to observe when specific large-effect restorers are already abundant in the population – as expected if CMS has been present for sufficient time.

As illustrated above, CMS is well understood theoretically and observed in plants but is exceedingly rare among animals (Vellnow et al., 2017). However, CMS has been recently reported in one animal: *Physa acuta*, a freshwater snail (David et al., 2022). In this species, two mitochondrial types conferring male sterility, D and K, have been discovered. They show extreme molecular divergence from one another, as well as from normal, fertile cytotypes (collectively called N), at all mitochondrial genes. The D mitochondrial type was discovered first, but evidence for corresponding nuclear restoration is still lacking (David et al. 2022). Recently however, the second CMS mitotype K revealed a different situation, with evidence that most K individuals captured in natural populations have a male-fertile phenotype, i.e., that their nuclear genomes contain genes restoring male fertility, while male-sterility reappears when the K mitotype is introgressed into a “naïve” laboratory genetic background maintained without contact with CMS (Laugier et al. 2024).

In this study, we explored the inheritance and specificity of restoration in *Physa acuta*. We aimed to characterize the variation in restoration for the K mitotype, specifically to test if the data are consistent with the existence of dominant-restorer alleles in natural populations where K is present. We also tested whether these restorers are specific to K or also have an effect on male-fertility of the D mitotype. Finally, we were interested in the determinism of male-fertility in our laboratory population, where the K-mitotype, although mostly male-sterile, still maintains a fraction of male-fertile individuals. This population may represent a truly “naive” state, i.e., the state of a population in which a new CMS factor would be introduced into the population’s background of standing variation; alternatively, it may have inherited, from many generations ago, the same restorer alleles as in natural populations where K is present, but these alleles are now at a low frequency.

To achieve these objectives, we phenotyped CMS K- and D-mitotype individuals with two different nuclear backgrounds (Lyon and Montpellier). The Lyon population has been in contact with both K and D (i.e., non-naive), while the Montpellier one has not (i.e., naive). To reveal genetic variation in restoration potential within the Lyon and Montpellier populations, we used two different approaches: we produced populations selected against or for restoration, and we analysed inbred lines. This combination of approaches enables us to explore restoration in various contexts and characterize it. Answers to these questions will provide insight into the CMS evolutionary dynamics and, for the first time, explore genetic variation for CMS restoration in animals.

## Material and methods

### Study system

*Physa acuta* is a cosmopolitan, hermaphroditic, predominantly outcrossing, freshwater snail, very common in natural freshwaters worldwide, and easy to raise in the laboratory. The generation time is about 6-8 weeks in the laboratory at 25°C; we maintain this snail in standard conditions (25°C, 12/12 photoperiod) using ground water changed at least once a week, and grinded boiled lettuce ad libitum as food. Cytoplasmic male sterility was first described in this species by David et al. (2022) and later completed by Laugier et al. (2024). Briefly two very divergent mitogenomes, named D and K, have been found to coexist with the normal (N) type in wild populations near Lyon (France). While N individuals are regular hermaphrodites, D and K are associated with male-sterile phenotypes, unable to play the male role in sexual interactions and to sire offspring. Nearly all the D (92%, David et al. 2022) individuals are male-sterile. In contrast, nuclear backgrounds restoring male-fertility of K snails are found in the Lyon populations where K is present. However, the same K mitotype, when introgressed into the nuclear background of a population from Montpellier (hereafter « naive », maintained for >80 generations in the laboratory without contact with CMS), produces around 70% male-sterile phenotypes (Laugier et al. 2024). The nature of the ∼30% residual fertility is still hypothetical and it is possible either that restorer genes are segregating at low frequency in the naive population (despite the absence of K mitotypes); or that the penetrance of CMS is incomplete, leaving a fraction of individuals with nonzero male fitness.

### General structure of the experiments

Our objective was to characterize, within both the naive (Montpellier) and the non-naive (Lyon) backgrounds, genetic variation for the potential to restore male fertility in CMS mitotypes. We did this in two ways (i) first, we tested whether we could select for high versus low restoration potential within the naive population (hereafter, “selected lines” dataset) (ii) second, we produced inbred lines from both populations and characterized genetic variance in restoration potential among lines, specially aiming to identify lines fixed for either restorer or maintainer (i.e., non-restorer) alleles (hereafter, “inbred lines” dataset). In both cases, lines were of the N mitotype, and their restoration potential was evaluated by crossing them with either a K or a D snail and then evaluating the male fertility of the offspring. In these crosses, the paternal strains were N while the maternal ones were K or D. As mentioned above, all (or nearly all) our D individuals are non-restored and mostly male-sterile (thus termed DS). However, two K strains are available, one with the original background from Lyon (with high restoration potential and mostly male-fertile, termed KF) and one with the naive background from Montpellier (low restoration potential, mostly male-sterile, termed KS). We used both fertile and sterile strains (K and D) as mothers in order to observe the effects of potential restorer alleles provided by the paternal strain, either in a predominantly homozygous (cross with the high restoration potential KF line) or in heterozygous state (cross with the low restoration potential KS and DS lines).

### Source populations and their history in the laboratory

The Montpellier population was founded in 2007 by mixing ten natural populations sampled around Montpellier, and maintained in the laboratory, in high numbers, since then (Figure 1). A spontaneous mutation in this population enabled the separation of an albino stock from the main population within the first few laboratory generations (see Noël et al., 2016). It has been maintained as an autonomous large outbred population in aquaria to this day (estimated age in 2023: 96 generations) and is entirely of the N mitotype and male-fertile. Albinism is due to a recessive allele, and albino mothers produce pigmented offspring when inseminated by a wild type snail. More recently, a new type of albinotic mutants appeared within the Montpellier stock, with a slightly different phenotype (see S1, Figure S1A). After some crosses (see S1, Figure S1B) we were able to establish it as an independent outbred stock (called alb-2) that complements with the first albino stock (now called alb-1): alb-1 and alb-2 are mutated at different loci, and crosses between them produce 100% pigmented individuals, as well as crosses between either albino type and pigmented homozygous snails.

**Figure 1:**
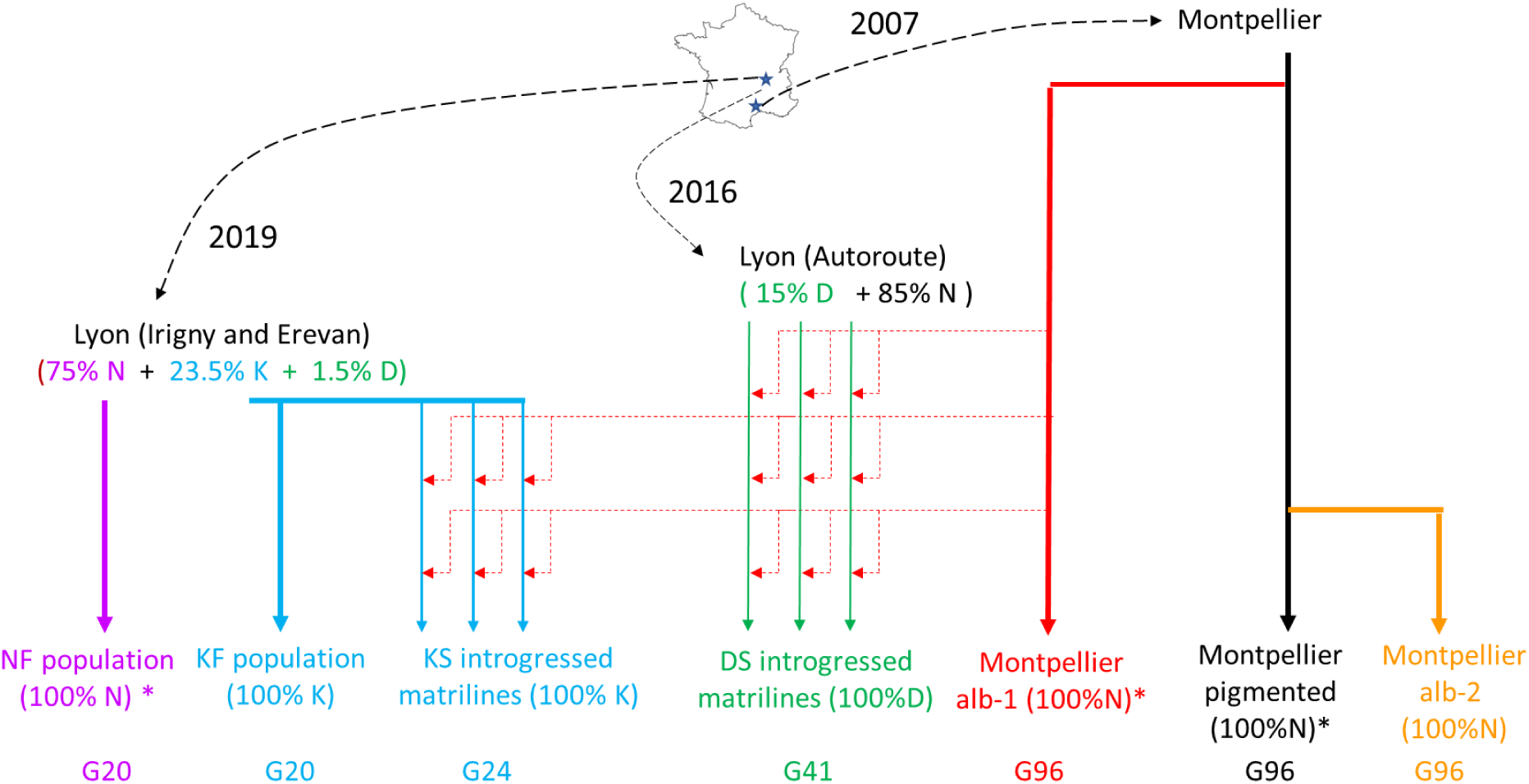
Source populations and their history in the laboratory since extraction from natural populations (indicated by dashed arrows, with year of sampling) Thick lines indicate propagation in large outbred populations in aquaria. Thin lines indicate propagation in matrilines, with insemination at each generation by sperm donors from another population (dotted arrows). The percentages of each mitotype (K, D or N) are indicated for the natural populations at sampling (from David et al.2022, Laugier et al. 2024) and for populations kept in the laboratory. The number of generations since extraction from natural populations in late 2022 – early 2023 (when most of the experiment took place) is indicated as G_XX_ below each stock. The numbers of independent matrilines are 12 for KS and 11 for DS. Populations from which inbred lines were extracted are indicated by a star (*). The coordinates of the populations of origin can be found in Noël et al. 2016 (Montpellier), David et al. 2022 (Lyon, Autoroute), and Laugier et al. 2024 (Lyon, Irigny and Erevan)

The Lyon populations were founded from two populations sampled in 2019 near Lyon mixed together (Irigny and Erevan, see Laugier et al. 2024). They initially contained a mix of K and N mitotypes, the former with mostly male-fertile phenotypes (see above) and all pigmented (wild-type). After mass-mating, we collected progenies from individual mothers that were typed by PCR as either K or N; and the K and N progenies were pooled into two separate tanks, to initiate pure-K and pure-N stocks (Figure 1). Each stock was kept as a separate population (hereafter KF and NF populations) and maintained as a large outbred population to this day (in 2023, KF and NF were at generation 20 after foundation). Each of the outbred populations (alb-1, alb-2, NF, KF) was maintained throughout as a metapopulation of 5 aquaria, each with 100-150 snails at reproductive stage, freely mating (« mass-mating »), with regular exchange of ∼10% individuals among aquaria; the eggs are separated from the parents for hatching as soon as there are enough on the aquarium walls, and a new cohort is started. Aquaria are fed once to twice a week depending on their life stage.

A final source of snails for our experiments is a set of lines in which the D or the K mitotypes have been introgressed into the naive Montpellier alb-1 background (David et al 2022, Laugier et al 2024). Briefly this introgression involves sets of « matrilines » each founded by a D or K ancestor from Lyon (wild population) and propagated by maternal descent. At each generation, each snail from a matriline is inseminated by an individual taken at random from the alb-1 stock to progressively replace the nuclear genome of the founder by the alb-1 background. At the time of the experiment the D matrilines were at the 41st generation and the K matrilines at the 20th generation. Their nuclear genome can be considered as totally replaced. Matrilines are maintained with pigmented phenotype and heterozygous at the alb-1 pigmentation locus to distinguish them from their albino mates: at each generation we obtain a mix of albino and pigmented progeny, and we keep only the latter to propagate the matriline. For the experiment, however, we could only use the albino progeny. Both K and D matrilines are mostly male-sterile (as mentioned above); for this reason we will refer to them as KS and DS respectively (by contrast, KF snails are restored and mostly male-fertile).

### Selection for high and low frequency of restoration of the K mitotype

The aim was to test whether there was variation from which restoration potential could be rapidly selected when a CMS mutant appears in a naive population. This was envisaged for the K mitotype because the penetrance of the male-sterile phenotype in KS individuals (mitotype K introgressed into the naive alb-1 background) is around 60-70%; if the residual fertility is genetic, it should increase or decrease under selection. The residual fertility of DS snails is too low to start a similar selection experiment.

The selection protocols are detailed in supplementary note S2. In short, to select against restoration, we kept the (N-mitotype) alb-1 sperm donors that, when crossed with a KS snail, produced only male-sterile K daughters. The selected sperm donors were then grouped and constituted the first generation of selection for low-frequency-of-restoration (LFR). We re-used offspring of these individuals as sperm donors for a second round of selection. Therefore, the LFR individuals used here are considered to be at their second generation of selection against restoration. The selection for high frequency of restoration (HFR) is simpler because selecting for fertility is easier than for sterility: we simply kept the offspring sired by male-fertile KS individuals each generation. The HFR population was selected for two successive generations, the same as the LFR. In the simplest scenario (fully penetrant, monodominant restorers), this selection should produce very contrasted restorer frequencies between LFR and HFR (0.035 versus 0.521, details in S2).

### Constitution of inbred lines

Inbred lines allow in principle to fix restoring and non-restoring genotypes as homozygotes. We created inbred lines from N-mitotype ancestors originating from either the Lyon or the Montpellier laboratory populations (more lines were made from Montpellier where restoration appears to be rare). Each ancestor produced a line by at least two generations of selfing (enforced by letting a virgin snail reproduce in isolation) and single-individual descent. Lines are not fully homozygous at the second generation (¼ of the initial heterozygosity is expected to remain) but the design is a compromise between the time required and the potential loss of lines by inbreeding depression over generations. In this experiment we used 9 lines from Lyon and 40 from Montpellier.

### Crosses

In total we tested the restoration potential of focal individuals from two selection lines (HFR and LFR) and from 49 inbred lines of two origins: Montpellier (naive population) and Lyon (non-naive population). To that end focal individuals were used to inseminate three types of individuals: DS, KS and KF. The resulting progeny have the focal individual as father and DS, KS or KF mothers. In each progeny we grew one to three G1 offspring until maturity and tested them for male fertility. If the father transmits restorer alleles, these alleles are present in the tested offspring either in a mostly heterozygous form (mothers DS or KS, with absent or rare restoration), or in a mostly homozygous form (mothers KF, with a high rate of restoration). The total number of progenies tested was 1508, as listed in Table 1.

### Assessment of male sterility

The assessment of male sterility of an individual is made by pairing it to a virgin partner for three days, letting the latter lay for three days in isolation, and observing the progeny. The studied individual is said to be male-sterile if either the partner does not lay any egg (failure to stimulate egg-laying, which is a normal outcome of insemination) or it lays eggs but the juveniles are 100% self-fertilized. Partners of male-fertile individuals in this situation normally produce several tens of eggs and juveniles are nearly all outcrossed. A small number of individuals had an intermediate status: outcrossed individuals were obtained but in abnormally low numbers (<10); in this case we recorded them as semi-sterile. In the analyses we grouped these together with fully male-sterile individuals (without appreciable consequence on the results). It is, in any case, important to realize that the male-fertility test is not perfect: even male-fertile individuals (with fertile mitotype) occasionally fail to inseminate their partner during pairing, with a probability <10% in our lab conditions.

We used body pigmentation to recognise outcrossed offspring from self-fertilized ones. Depending on the paternal pigmentation genotype, we used either albino alb-1 or alb-2 partners in such a way that any pigmented offspring they produced must have been sired by the tested individuals. We therefore simply counted the number of pigmented offspring in the progeny after incubating clutches for 13 days (we stopped counting at 20 and noted 20+ if all offspring were pigmented, and counted exhaustively all pigmented and albino offspring otherwise, including those that were well-developed though not yet hatched, if any). In some cases (heterozygous individuals tested with an alb-1 partner) outcrossed offspring were expected to be 50% pigmented, 50% albino instead of 100% pigmented; in this case we set the threshold for semi-sterility to 5 pigmented (i.e. the expectation for 10 outcrossed offspring) instead of 10 (see above).

We tested male fertility in G1 individuals and in their DS, KS, KF mothers too (see supplemental figure S3). Many of the latter were pigmented and mated with albino N-mitotype fathers; thus, the father could act as a partner for testing the male-sterility of the mother, and we kept the father’s progeny to that end. In case the father was a pigmented individual and/or the mother an albino, this was not possible so we provided a second partner to the mother, after she had laid eggs, and this second partner was alb-2 so that outcrossed offspring would be recognised (alb-1 and alb-2 are complementary lines and produce pigmented babies upon cross-fertilization). Because of the size of the experiment, it was spread over nearly two years from 2022 to early 2024, with four phases treated as temporal blocks.

### Statistical analyses

We analysed separately the differences between the selected lines (HFR and LFR) on one hand, and the variation among inbred lines on the other hand. The male-fertility status of the offspring of selected lines was treated by GLMMs as a binary 0/1 variable (Binomial distribution, logit link). The main model included the maternal type (KF, KS or DS), the selected line (HFR or LFR) and their interaction as fixed effects, and the identity of the mother and the block as random effects. We also made more detailed models including the maternal phenotype (male-fertile or male-sterile) in the fixed effects. The male-fertility status of the offspring of inbred lines was treated in a similar way, except that the fixed effects included: the geographic origin of the inbred line (Montpellier versus Lyon), the maternal type (KF, KS or DS) and their interaction; and the random effects included the identity of the inbred line, in interaction with maternal type, in addition to block and mother identity. In the inbred lines dataset, we also used the model estimates of variance among paternal lines to estimate heritability under the quantitative-genetic threshold model, as proposed by Ehlers et al. 2005. The computation procedure, based on (de Villemereuil et al., 2016), is detailed in supplement S5. All tests were performed by likelihood-ratio chi-squares, comparing models with and without the effect of interest with all other effects kept. The models were run using the lme4 package in R (Bates et al. 2015).

### Predictions

The selected lines inform on the nature of the variation in male-fertility observed when CMS is introduced into a naive background (the Montpellier laboratory population), i.e., why is there a residual male fertility in K individuals introgressed into this background? If there exist restorer alleles with large effects, the frequency of male-fertility of K individuals sired by HFR fathers should exceed that of their congeners with LFR fathers. Depending on dominance, this effect should be more visible in different circumstances. A dominant allele transmitted by fathers makes a difference when the mother does not herself bring the same allele (in our case, with male-sterile mothers, and/or with mothers from the KS strain). A recessive allele, on the contrary, would have more effects when the mother is male-fertile and/or from the KF strain. Finally, the specificity of restoration with respect to the mitotype can be tested by checking whether the LFR/HFR difference, if any, affects male-fertility when the mother is of the D mitotype.

The inbred lines allow to document the nature and specificity of restoration in both the Montpellier and Lyon populations. Based on previous studies (Laugier et al. 2024) we expect that K-mitotype individuals sired by Lyon lines (a largely restored population) will have more male-fertile offspring than those sired by Montpellier. We expect, if restoration has a simple genetic basis, that some lines will be homozygous for restorers and others for maintainer alleles, generating high variation in male-fertility among lines, either when crossed with KS / male-sterile mothers (if restorers are dominant) or when crossed with KF / male-fertile mothers (if recessive). Similar effects could be observed upon crossing with DS if there exist restorers efficient against CMS brought by the D mitotype; restoration may be specific or not (in the latter case, the same lines should restore fertility in both D and K).

## Results and Discussion

Our results indicate that male-fertility in *Physa acuta* is influenced by two distinct systems. In the Montpellier background the penetrance of male sterility induced by the CMS mitotypes is always high but not 100%, and has significant but limited heritability. In contrast, our findings suggest that in the Lyon population, one or a few dominant restorer alleles inducing near-complete male-fertility in the K mitotype are present at high frequencies.

### Quantitative modifiers of male-sterility penetrance in the Montpellier background

All crosses involving KS or DS mothers with fathers from either inbred or selected (HFR/LFR) lines had a 100% Montpellier background. In these crosses, maternal phenotypes (Figure 2) were mostly male-sterile, as expected based on previous studies (see Materials and Methods): we found only 5.7% of male-fertile individuals among DS mothers and 27.2% among KS ones. Similar proportions were obtained in their offspring, when the father also came from Montpellier (HFR or LFR lines, Figure 3, Montpellier inbred lines, Figure 4; compare with Figure 2). Accordingly maternal type had a highly significant effect in both datasets (selected lines and inbred lines) and in both generations (mothers and offspring, Figure 2, Table 2 and 3).

**Figure 2:**
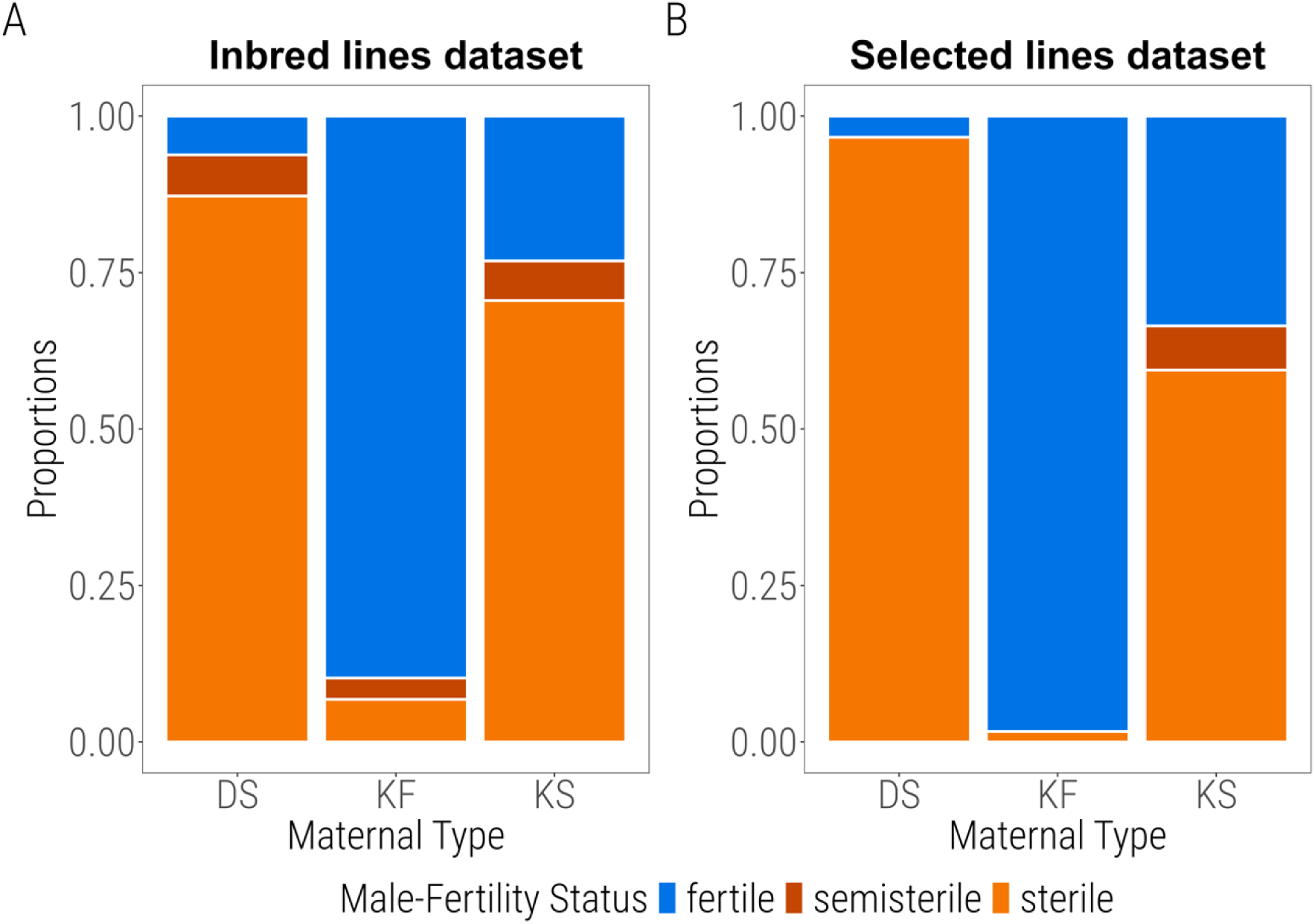
Proportions of individuals with male-sterile, male-fertile and male-semi-sterile phenotype in the maternal generation. Proportion are represented for both datasets (A: Inbred lines, B: Selected lines) as a function of maternal type (DS: D-mitotype introgressed into the alb-1 background from Montpellier; KF: K-mitotype within its native background from Lyon); KS : K-mitotype introgressed into the alb-1 background from Montpellier. The two datasets are presented separately because they were made at different times. In both datasets the proportion of male-fertile individuals differed very significantly among types (Binomial GLMMs, χ²_1df_ =119.9, and 36.6 respectively for the inbred lines and selected lines datasets, both *p* < 0.001).

**Figure 3:**
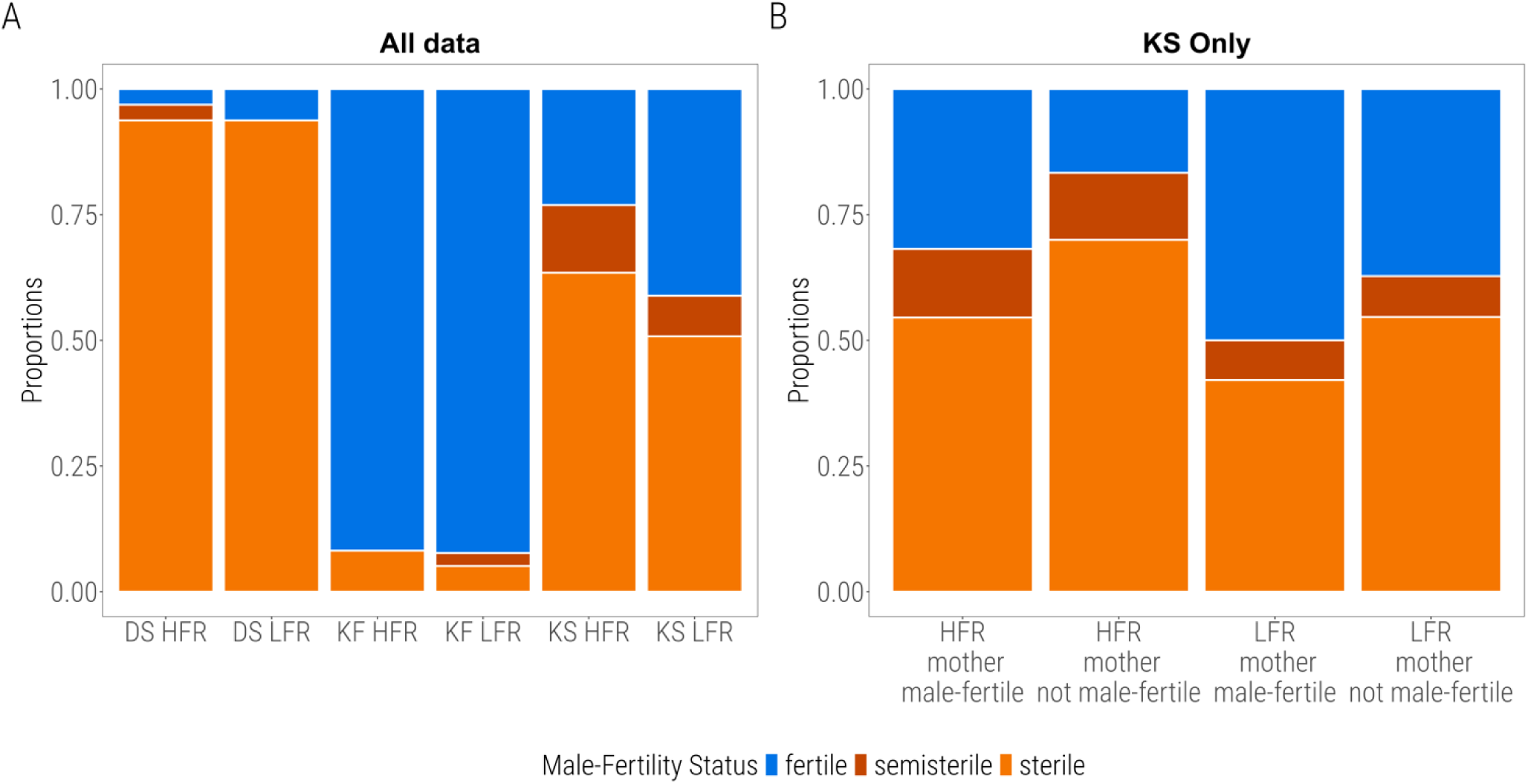
Proportions of individuals with male-sterile, male-fertile and male-semi-sterile phenotype, selected lines dataset. A: proportions of male-fertile and male-sterile phenotypes as a function of paternal selection line (HFR versus LFR), maternal type (DS, KF, KS). B: detailed view of proportions in offspring of KS mothers, as a function of paternal line and maternal phenotype (male-fertile versus not male-fertile). GLMMs indicate that both the paternal line (χ²_1df_ = 4.65, p = 0.031); and the maternal phenotype (χ²_1df_ = 4.48, P = 0.034) are significant but not their interaction (χ²_1df_ = 0.15, p = 0.70). Similar divisions were not considered in the DS and KF category because of the low numbers of male-fertile and non male-fertile mothers respectively.

**Figure 4:**
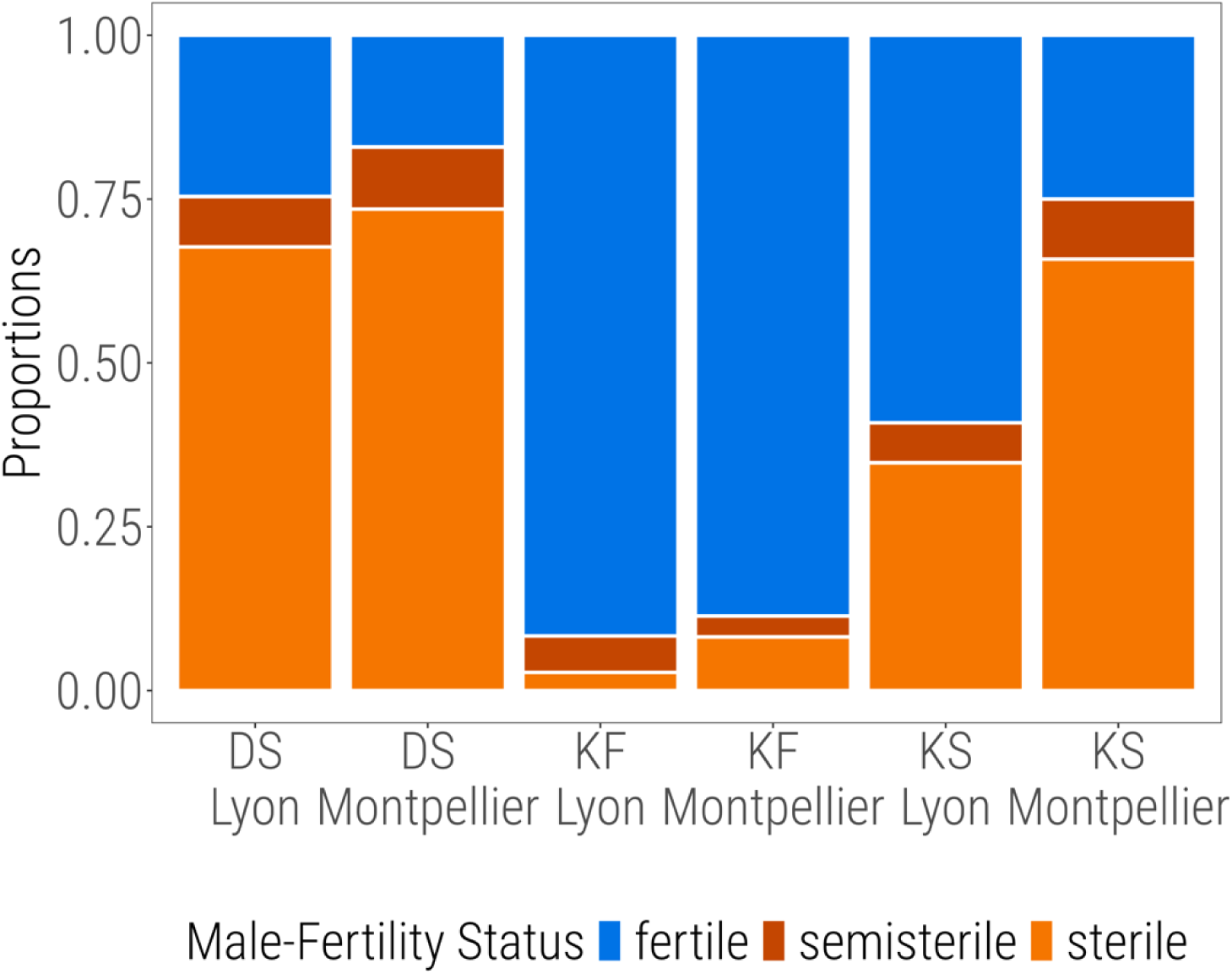
Proportions of male-fertile and male-sterile phenotypes as a function of the origin of the paternal inbred line (Lyon or Montpellier) and the maternal type (DS, KF, KS).

However, within the KS maternal type, the male-sterile phenotype was not faithfully transmitted from mother to offspring. Indeed, the maternal phenotype had no significant effect in the inbred lines dataset (χ²_1df_ = 0.018, p = 0.895, supplemental figure S4A), and in the selected lines dataset male-fertile mothers produced only slightly more male-fertile daughters than male-sterile mothers did (Figure 3, χ²_1df_ = 4.48, p = 0.034). Similarly, in the selected lines experiment, fathers from the two selected populations (HFR and LFR) slightly differed in restoration potential (p = 0.039, Table 2). However, the difference was small and opposite to our expectations, as the LFR line—selected for low restoration frequency—produced more male-fertile offspring than the HFR line (Figure 3). This suggests that selection on the HFR and LFR lines was ineffective, and that differences between them reflect genetic drift. The Montpellier background is therefore devoid of dominant alleles fully restoring male-fertility in the K mitotype, as such alleles would result in strong differences in male-fertility between LFR and HFR in the direction of selection (3.7% versus 52%, see supplement S2). Genetic variation is rather expressed as a quantitative variance in the penetrance of male-sterility (rather than a binary 100% sterile vs. 100% fertile outcome). In such conditions, male fertility of an individual is an imprecise indicator of its genetic potential, reducing the effectiveness of selection relative to drift, particularly given the small number of individuals meeting selection criteria. Results of the inbred lines experiment are also in agreement with this hypothesis: in the KS context, frequencies of male-fertile phenotypes significantly differ among paternal inbred lines from Montpellier (Figure 5; Table 3; *h*²=0.29 under the threshold model, table S5) but remain on average very low and below 50%.

**Figure 5:**
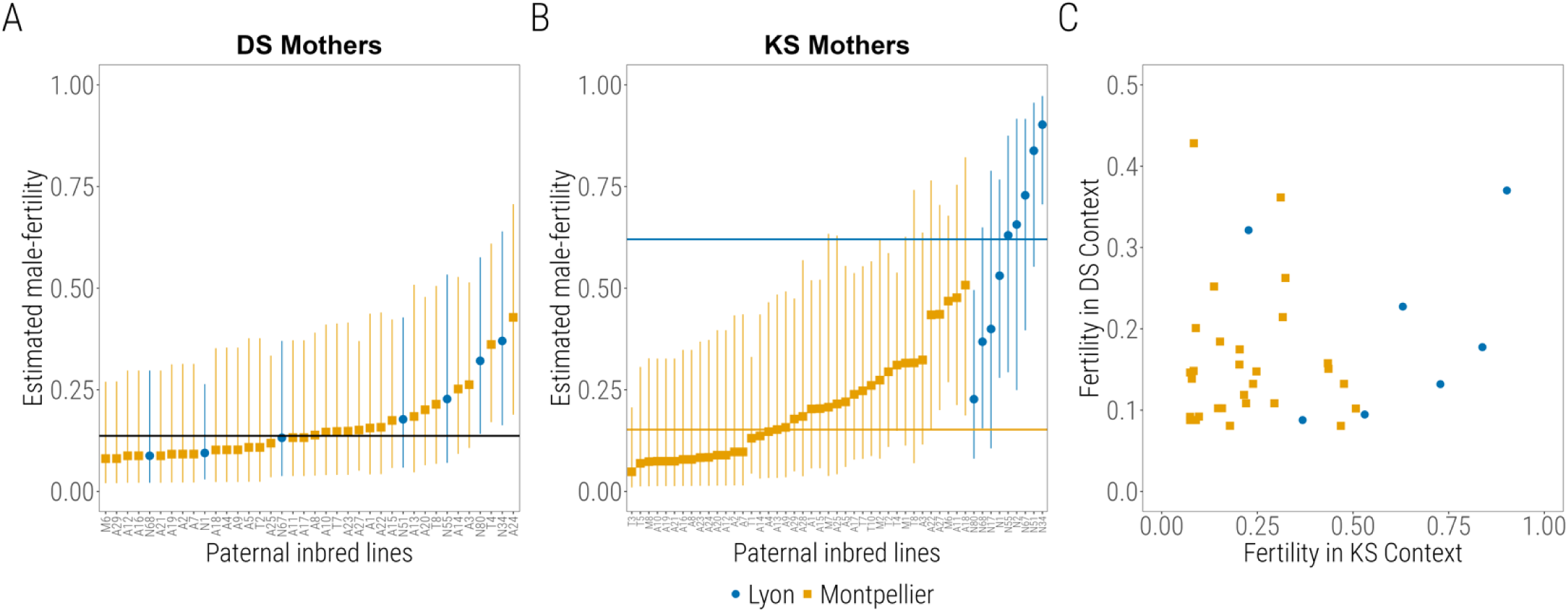
Proportions of male-fertile individuals in offspring sired by various inbred lines from Montpellier and Lyon, in a DS maternal context (A) or KS maternal context (B) The estimates are extracted from binomial GLMMs (BLUPs), de-logited to put them back into natural scale, and presented with 95% confidence intervals. The lines are arranged by increasing male-fertility, and separated by origin in the KS context, as the population of origin has a significant effect in this context (in DS context, this is not the case). Horizontal lines provide mean proportions of male-fertile individuals estimated by the model (de-logited intercepts), over all paternal lines (DS context) or separately for each population of origin (KS context). Note that there are more paternal lines tested in KS than in DS context. C: the performance of lines tested in the two contexts are plotted together (each dot is a paternal inbred line). The correlation is not significant (Spearman’s ρ = 0.22, *p* = 0.184).

**Table 3:** Generalised linear mixed models on the binomial male-fertility variable (fertile versus sterile or semi-sterile), inbred lines dataset. Legend as in Table 2. The block random factor was omitted because the estimated variance was 0.00. ^(1)^ the test gives the significance of the variance explained by the paternal line, in interaction with maternal type. Because of the significant interaction, the variances (in logit scale) explained by paternal line are examined within each maternal type separately. In each case we also provide separate estimates of variance for the inbred lines derived from Montpellier and Lyon populations.

| Data |  | Fixed effects |  |  | Random effects |  |
| --- | --- | --- | --- | --- | --- | --- |
|  | maternal<br>type (DS,<br>KF, KS) | paternal origin<br>(Lyon,<br>Montpellier) | interaction | maternal ID | paternal line |  |
| all | $\chi^2_{df2} = 80.61$<br>$p < \mathbf{10^{-15}}$ | $\chi^2_{df1} = 9.73$<br>$p = \mathbf{0.002}$ | $\chi^2_{df2} = 5.37$<br>$p = 0.068$ | $\chi^2_{df1} = 7.16$<br>$v = 0.92$<br>$p = \mathbf{0.0037}$ | $\chi^2_{df5} = 16.75$<br>$p = \mathbf{0.005}^{(1)}$ | |
| only<br>DS | | $\chi^2_{df1} = 1.439$<br>$p = 0.203$ | | $\chi^2_{df1} = 0.18$<br>$v = 0.29$<br>$p = 0.387$ | $\chi^2_{df1} = 6.58, v = 0.80$<br>$p = \mathbf{0.005}$<br>(in Montp: $v = 0.77, p = \mathbf{0.011}$,<br>in Lyon: $v = 0.79, p = 0.08$) | |
| only<br>KF | | $\chi^2_{df1} = 0.28,$<br>$p = 0.597$ | | $\chi^2_{df1} = 0.89$<br>$v = 1.32$<br>$p = 0.174$ | $\chi^2_{df1} = 0.00, v = 0.00, p = 0.5$<br>(in Montp: $v = 0.00,$<br>in Lyon: $v = 0.00$) | |
| only<br>KS | | $\chi^2_{df1} = 14.84,$<br>$p = \mathbf{0.0001}$ | | $\chi^2_{df1} = 6.54,$<br>$v = 1.00,$<br>$p = \mathbf{0.005}$ | $\chi^2_{df1} = 32.4, v = 1.58,$<br>$p = \mathbf{6.2 \cdot 10^{-9}}$<br>(in Montp: $v = 1.45, p = \mathbf{0.0005}$<br>in Lyon: $v = 1.97, p = \mathbf{8.5 \cdot 10^{-5}}$) | |

Within the DS maternal type, numbers of male-fertile phenotypes in mothers and offspring are too low to test mother-offspring resemblance, and to select for or against restoration. However, the effect of paternal lines in DS context are similar to observations in KS context: although they differ significantly in their restoration potential (Figure 5; Table 3, *h*² = 0.168 under the threshold model, Table S5), none of them restores more than 50% male-fertility. In addition, proportions of male-fertility in paternal lines are not significantly correlated between the DS and KS contexts (Figure 5).

From these results, we conclude that (i) in nuclear genomes from Montpellier we did not detect strong-effect, dominant alleles fully restoring male fertility in K and D mitotypes; rather the observed fractions of male-fertile phenotypes reflect an incomplete penetrance of CMS, (ii) penetrances of K-induced and D-induced male sterility behave as uncorrelated quantitative traits with low genetic variance. This aligns with the threshold trait model (Ehlers et al., 2005), where the same genotype can be either male-fertile or male-sterile with a genetically determined probability. Additionally, we observed some cases of partial male sterility (“semi-steriles”; Figures 2, 3, 4), though their low numbers precluded separate analysis. This suggests that a binary classification of male sterility simplifies the actual phenotypes. In polygenic restoration systems, full restoration of CMS may be rare, with male fitness varying quantitatively among restored CMS hermaphrodites (e.g., Dufay et al., 2008). Our findings align with previous studies in non-crop species that rejected simple Mendelian models of restoration (e.g., *Thymus vulgaris*, Belhassen et al., 1991; Charlesworth & Laporte, 1998; *Lobelia siphilitica*, Dudle et al., 2001).

### Strong-effect, K-specific dominant restorers in the Lyon background

Crosses involving KF mothers, and/or fathers from the Lyon population (i. e., parents with Lyon background) revealed that male fertility of the K mitotype has a distinct genetic basis in Lyon (where K naturally occurs) compared to Montpellier. First, KF mothers were nearly all male-fertile (92.3%, Figure 2), and consistently produced male-fertile daughters, with no effect of paternal origin - including when the father was from a restorer-free Montpellier background (HFR, LFR, Montpellier inbred lines, Tables 2 and 3, Figures 3 and 4). This suggests that KF mothers transmit dominant restorer alleles. Second, fathers from Lyon produced on average 62% male-fertile offspring when crossed with KS mothers. Thus, they significantly differ from fathers from Montpellier (p =0.0001, Table 3, Figure 4). In the KS context, variance in male-fertility among paternal lines from Lyon was highly significant and higher than in Montpellier (*p* = 8.5 10^-5^, Table 3; *h*² = 0.40, Table S5). Paternal lines from Lyon spanned a range from ∼25 to 95% male fertility, with two-thirds of the lines above 50%. Interestingly, the percentages were continuously distributed in this range (Figure 4 and see supplemental figure S4B), not bimodal, which suggests the involvement of more than one locus – F2 or backcrosses could be made in the future to explore this hypothesis.

In summary, genes from Lyon, transmitted by both mothers and fathers, are able to restore high frequencies of male fertility in presence of a K mitogenome, even when put in heterozygous state with a (restorer-free) hemigenome from Montpellier. Note however that Montpellier-Lyon crosses yielded higher frequencies of male-fertility when the parent from Lyon was the mother (KF mothers x HFR or LFR fathers) than when it was the father (KS mothers x fathers from Lyon inbred lines). This likely reflects differences in recent histories in the laboratory. KF mothers originated from a population maintained at 100% K mitotype for 20 generations, where selection for restorer alleles has probably been strong. Conversely, Lyon inbred lines derived from a 100% N population (see Figure 1), where restorers were functionally silent and could have declined if costly. Further replication is needed to confirm this hypothesis.

### Restoration is specific to the K mitotype

Overall, most of the D-mitotype individuals were male-sterile irrespective of the nuclear background. Unlike in the KS context, we found no evidence for dominant restorers. None of the paternal lines (even those that restore near 100% fertility in K context) produced more than 50% male fertility, although we observed weak among-line variance for penetrance (h² = 0.21, table S5). Because the Montpellier population, unlike Lyon, has had no recent contact with the D mitotype, we would expect a higher rate of male-fertility in Lyon if selection has acted on this variation. However, we found no significant difference (p = 0.203, Table 3) and the distribution spans the 0-50% interval with an apparently random distribution of lines from Montpellier and Lyon (Figure 4). Thus, strong-effect dominant alleles restoring male fertility in K context, present in the Lyon population, do not have the same effect on mitotype D.

Why did we find K-specific restorers but not D-specific ones? In the wild the D mitotype was found only at low frequency (14% or less, David et al., 2022; Laugier et al., 2024). Thus, D may not be frequent enough for a restorer to be selected. The high evolutionary rate detected in the D mitotype (David et al. 2022) could result from an arms race with restorer genes that are now extinct or rare or have lost their ability to restore fertility following a recent evolution of the D mitochondrial genome itself. According to the frequency-dependent model, restorers should spread rapidly and, if they have no or little fitness cost, can go to fixation. However, nuclear restorers are expected to be rare if they have a high cost (Gouyon et al., 1991; Dufaÿ et al., 2007). Founder effects causing a reduction of the nuclear diversity could also lead to the local absence of nuclear restorers in recently founded populations (e.g. Manicacci et al., 1997; Laporte et al., 2001). Therefore, we can hypothesize that the evolution of restorers specific to the D mitotype might be constrained by a high cost or founder effects.

### Two stages in the maintenance of cyto-nuclear polymorphism

Most models have studied the maintenance of the cytoplasmic and nuclear genetic variation in gynodioecious populations assuming strong and dominant restorer alleles at a few loci. Such alleles are indeed the most likely to be under strong selection and undergo selective sweeps in populations invaded by new CMS genes. Depending on their associated costs, they may either reach fixation or stabilize at intermediate or cyclically variable frequencies together with their corresponding CMS cytotypes (Gouyon et al., 1991; Dufaÿ et al., 2007). Gynodioecious populations observed today are mostly in the polymorphic state and the restorers found indeed include large-effect alleles with dominant action (e.g. de Haan et al., 1997; Dufay et al., 2008, 2009). In addition, this action is usually specific to each cytotype, revealing either an arms-race coevolutionary history specific to each cytotype-restorer pair, and/or the fact that each new CMS cytotype has a mode of action that differs so strongly from previous ones that existing restorers are unlikely to neutralize it. In *Physa acuta,* the K mitotype, and the restorers present in the natural population where it was found, are likely to be in this situation (alleles with strong, dominant, and cytotype-specific effects).

While such situations are expected to be frequent in stable gynodioecious populations, they may not represent the initial stages when a new CMS mutant arises in a ‘naïve’ population, where nuclear genes have not yet undergone selection to suppress its effects. According to Lewis (1941), the spread of CMS in the population will in this case become limited only by pollen limitation when females become so abundant that many of their ovules remain unfertilized. Our results suggest however a way by which such populations may lower the impacts of pollen (or, in the case of *Physa*, sperm) limitation: CMS may not be fully penetrant, leaving a residual production of male gametes sufficient to fertilize the eggs of mates when there are no normal male-fertile congeners. Even in a naïve population, pre-existing standing variation may include variants that happen to modulate this production. This probably corresponds to the situation of the K-mitotype when introgressed into the “naïve” Montpellier population, and of the D mitotype in both the Montpellier and Lyon populations. Here, we mean by “naïve” that it has either never had any history of selection in presence of the K and D mitotypes, or that the contact is too recent for large-effects restorers to have emerged, or too ancient for them to have persisted if they ever existed. The background variation acting on the penetrance of CMS could produce a reservoir of variation within the population, by which male fertility can start to increase, until the appearance of proper restorers.

The emergence of restoration could therefore often occur in two stages: an initial stage involving predominantly quantitative, weak modifiers of CMS penetrance, followed by a second stage in which dominant, strong restorer alleles may emerge and be selected at one or more loci. This process could ultimately lead to either fixation or mixed determinism of male fertility. While this interpretation is consistent with our findings, further investigation would be required to confirm its generality. Although theoretical models generally consider a simple genetic determinism for restoration (but see Frank, 1989; Bailey & Delph, 2007), empirical studies on wild plants have often rejected this assumption as mentioned above (e.g. Charlesworth & Laporte, 1998; Koelewijn, 2003). A complex determinism of restoration implies that within a population, CMS individuals may be partially restored (i. e. male fertility may be more of a continuous than binary 0/1 distribution) and this could affect the degree of selection for restorer alleles. Then, the positive selection of restorers could slow down and ultimately modify the conditions of maintenance of cytonuclear polymorphism, favouring the maintenance of females in gynodioecious populations (Bailey & Delph, 2007).

## Conclusion

This study shows that as in plants, the genetic determinism of male-sterility in *Physa acuta* is complex and suggests that it encompasses two different layers: one of weak, quantitative modifiers of the penetrance, not necessarily coevolved with CMS, the other of strong-effect restorer alleles with dominant action, that were selected in the presence of CMS. Moreover, the way the phenotype is measured could influence our perception of the restoration potential. Indeed, the male fertility status was evaluated in the absence of competition; it will be interesting to test whether restoration appears efficient when male-male competition is allowed. Additionally, the sexual phenotypes associated with D or K can be influenced by temperature (Bererd et al., 2025), emphasizing environmental effects on male-sterility dynamics. The genetics of restoration may also influence the maintenance of cyto-nuclear polymorphism by acting on restoration costs. This could be evaluated by looking at pleiotropic effects of nuclear background with different restoration potentials on male and/or female fitness in non-CMS cytotypes, although plant studies suggest that such costs are difficult to detect (but see de Haan et al., 1997; Bailey, 2002; Dufay et al., 2008). Finally, identification of QTLs for male-fertility restoration would be necessary for a full understanding of its genetic architecture in different populations.

