## Supplementary file for "Complex genetic determinism of male-fertility restoration in the gynodioecious snail *Physa acuta*"

**Supplementary materials**

### S1: Origin and extraction of the alb-2 stock

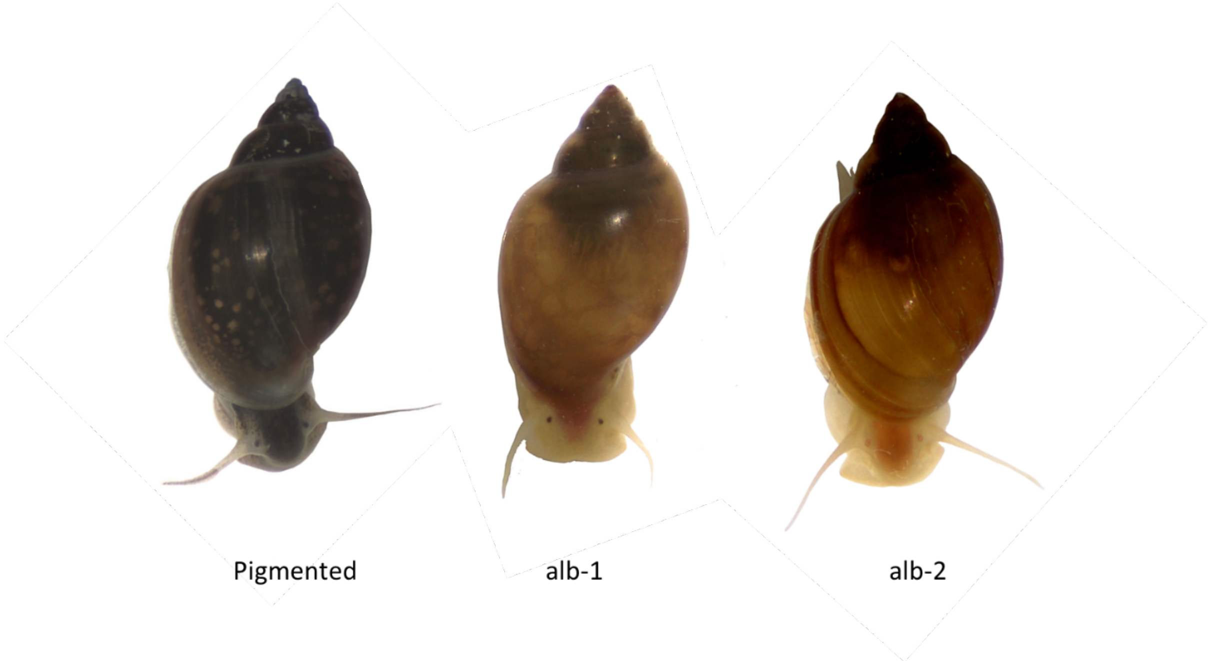


**Figure S1A: Phenotypes of three adults from the pigmented, alb-1 and alb-2 populations maintained in Montpellier.**

The alb-2 stock was initiated from a single individual with depigmented phenotype that appeared within a self-fertilized brood of a parent from the Montpellier pigmented stock, generation 74 in December 2020 (see Figure 1). We suspected it could be a different mutant from the alb-1 albino stock already available, because Dillon and Wethington (1992) mentioned the existence of two complementary albino strains in *Physa acuta* (*P. heterostropha* in their paper, but later synonymised). In addition, the adult phenotype of the new albino was slightly different from the alb-1: the dark pigments seemed completely absent, so that the body appeared slightly more orange (versus beige to yellowish for the alb-1), and the eyes were barely visible (Figure S1A). These traits are however very difficult to ascertain on hatchlings or juveniles because the eyes of alb-1 hatchlings are barely visible too.

We first let the newly discovered albino individual self-fertilize and obtained four $G_1$ offspring, each of which was crossed with a different individual belonging to the alb-1 population (complementation test, Fig S1B). All the $G_2$ obtained were pigmented, suggesting that the new type, now named alb-2, and the alb-1 were mutated on different genes and that both mutations were recessive to their respective wild-type pigmented allele. From the supposedly double-heterozygous $G_2$ we created a $G_3$ (avoiding crosses between full sibs) among which we selected only the non-pigmented phenotypes (which therefore were, according to our hypothesis, homozygous for albinism at least at one of the two loci; however we could not at that stage recognise different albino phenotypes from one another with certainty). Within the $G_3$ we wanted to select those that were homozygous for the albinism allele at the second locus, but homozygous for the wild-type allele at the first; i.e. individuals that, upon crossing with the alb-1 stock, would give 100% pigmented phenotypes. To that end we paired each of 155 $G_3$ with a different virgin DS individual of albino phenotype. The DS are male-sterile so they act as females in this cross, and their genetic background is that of the alb-1 population. We let the DS partners lay and in case the offspring were 100% pigmented, we selected the corresponding $G_3$ individual. We found 4 such individuals and left them together to mate; their albino offspring constituted the $G_4$. This $G_4$ was, given the low numbers of ancestors involved, rather inbred; to constitute a population with a larger genetic basis, we mated 22 of these $G_4$ each to a different pigmented individual from the Montpellier pigmented population. The $G_5$ obtained was 100% pigmented (as expected) and we retained 150 individuals that were left to mate in two large aquaria. We retrieved a large number of $G_6$ juveniles among which we selected 100 from the minority that had the alb-2 phenotype; these were used to found the alb-2 stock which has since then been kept as a large autonomous population (5 aquaria of ~100 individuals each with regular exchange of individuals among them). At the seventh generation we also crossed eight alb-2 each to a distinct alb-1 individual and checked that all offspring were pigmented as expected.

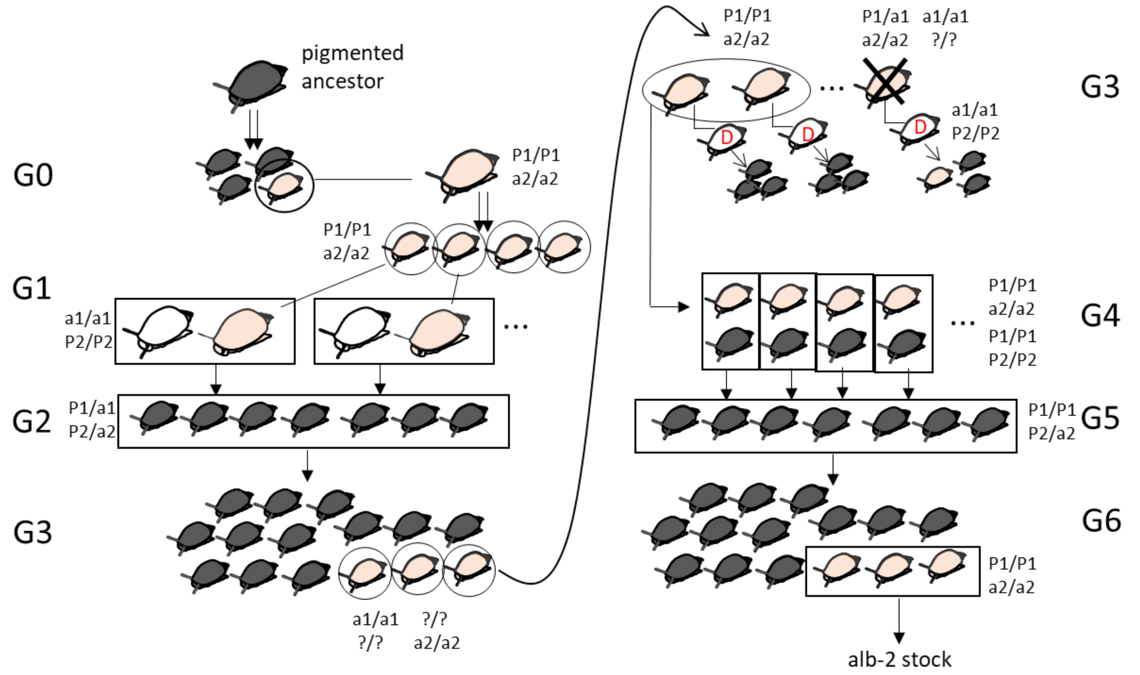


**Figure S1B: Derivation of the alb-2 stock.** Dark snails are of pigmented phenotype, light ones of albino phenotypes (the white and pink serve to distinguish snails but the two forms of albino phenotype were not recognised visually). The inferred genotypes at two loci (each with a pigmented dominant P; and an albino recessive a) are indicated but were not known, they were inferred a posteriori from the results of the crosses. $G_0$: an albino individual is identified in the self-fertilized brood of a pigmented ancestor from the Montpellier stock. $G_1$: the albino is replicated by selfing, each of four offspring is mated with an alb-1 individual. $G_2$: all offspring are pigmented and mated to one another. $G_3$: the albino phenotypes are selected and test-crosses made with D individuals with alb-1 background; $G_3$ that produce 100% pigmented offspring are selected (N = 4) and mass-mated together the others are eliminated. $G_4$: each of 22 offspring is mated with an individual of the Montpellier pigmented stock, to enlarge the genetic basis of the alb-2 population. $G_5$: all individuals are pigmented and left to mass-mate together (N=150). $G_6$: A minority of albino phenotypes are found; they are selected (N>150) to found the alb-2 population.

### S2: selection of the HFR and LFR lines

*Selection of the HFR (« high frequency of restoration ») line.*

The base population is the alb-1 Montpellier stock (with N mitotype), and the KS matrilines which have the same nuclear background as the alb-1 stock, except that the mitotype is K. Most KS individuals are male-sterile, however between 1/3 and ¼ of them are « male-fertile » (in the sense that they can fertilize a virgin N individual, and obtain >10 paternities this way).

The HFR selected line resulted from two successive crosses between K and N individuals as follows. In the first cross, multiple pair-crosses between KS pigmented and alb-1 individuals were made (e.g $K_1xN_1$, $K_2xN_2$, $K_3xN_3$ … $K_nxN_n$). Note that all the KS individuals used were daughters of male-sterile mothers. In each pair, the two partners were then isolated to lay separately. By phenotyping the progeny mothered by N individuals, we could assess male-fertility in its K mate (the K individual was said to be male-fertile if >5 pigmented babies were obtained from the N partner). The selection step to increase restoration frequency was as follows: If, according to the fertility test, the K individual of a pair was male-fertile, the offspring of both members of the pair were considered part of the first selected generation, otherwise they were discarded. In addition we kept only pigmented offspring of the K progeny, and albino offspring of the N progeny; the conserved offspring were raised to adulthood and used for a second cross. In this second cross, pairs with a K individual (pigmented) and a N individual (albino) were made again, taken from different families to avoid inbreeding. We again conserved only pairs for which the K individual turned out to be male-fertile, thus imposing a second selection step. In these pairs we conserved the N offspring only, and pooled them together to constitute de HFR stock.

Below(Table S2A) we give an expectation for the change of frequency of a restorer allele R during selection (noting m the corresponding maintainer allele), under the simplest scenario of monodominant restoration with full penetrance (i.e. RR and Rm individuals are male-fertile, and mm individuals male-sterile, in a K cytoplasm), noting S0, S1, S2 successive generations of selection. To simplify matters we assume that the starting frequency of R is 1/3 in the alb-1 source population (corresponding approximately to the observed frequency of male-fertile phenotypes in K offspring of male-sterile K mothers). In these conditions the frequency of R in the HFR population is expected to be 50/96 and the daughter of a male-sterile KS mother and HFR father should be male-sterile with probability 46/96 instead of 2/3.

**Table S2A: Expected frequencies in the HFR line, assuming monodominant restoration.**

| | K individuals | | | N individuals | | |
|---|---|---|---|---|---|---|
| Genotype | RR | Rm | mm | RR | Rm | mm |
| Gametic frequencies in alb-1 | | | | f(R)=1/3 | | f(m)=2/3 |
| Parents ($S_0$) before selection | 0 | 1/3 | 2/3 | 1/9 | 4/9 | 4/9 |
| $S_0$ after selection | 0 | 1 | 0 | 1/9 | 4/9 | 4/9 |
| $S_1$ before selection | 1/6 | 3/6 | 2/6 | 1/6 | 3/6 | 2/6 |
| $S_1$ after selection | 1/4 | 3/4 | 0 | 1/6 | 3/6 | 2/6 |
| $S_2$ | 25/96 | 50/96 | 21/96 | 25/96 | 50/96 | 21/96 |
| Gametic frequencies in HFR | | | | f(R)= 50/96 | | f(m)=46/96 |

*Selection of the LFR (« low frequency of restoration ») line.*

The LFR population derives from the same sources as the HFR and started with similar pair-crosses in the S0. This time, we conserved the N partner when its K mate was male-sterile i.e. the progeny of the N individual was either missing, or devoid of pigmented offspring ; in all cases we waited until the N individual self-fertilized. In parallel two pigmented babies of the male-sterile K mother were grown to adulthood and tested for their male sterility. If both were male-sterile (like their mother), suggesting that their father did not transmit them restoration genes, the self-fertilised progeny of their N Albino father was kept. On the other hand if either of them was male-fertile, the self-fertilised progeny was discarded. This whole protocol was repeated several times, adding the self-fertilized progenies each time to the same aquarium (approximately 15 progenies total) resulting in a self-sustained population called LFR 1 (Low Frequency of Restoration, 1st generation of selection). The population produced G1 individuals that we used as N partners for new KS individuals, to repeat the whole cycle of selection, yielding the LFR 2 population used in this study. The complexity of the selection step, requiring to eliminate most of the progenies, explains the low number of families that founded the LFR population so this population was exposed to important genetic drift compared to the ancestral alb-1 stock ; however the selection applied is very strong so it should be visible, if it applies to alleles with strong effects. For example, in the monodominant restoration scenario (see above) a *RR* parent is systematically eliminated by the selection process (because all his offspring are male-fertile) ; a *mm* parent is systematically kept (because all his offspring are male-sterile) and a heterozygous *Rm* parent is retained with probability ¼ (the probability that it gives the *m* allele to two independent offspring). To give a rough idea, if the frequency of R was initially 1/3, it should be 1/10 after the first generation of selection, and 0.037 after two generations (recall that the corresponding expectation is 50/96=0.521 in the HFR line).

**S3: Assessment of male sterility in G1 individuals and their mothers.**

For clarity, only one example is shown, where the mother of individual G1 is a DS individual. However, the same protocol applied to KS and KF mothers. **A:** Individual G1 had an N alb-1 father and a pigmented DS mother. To assess the male fertility of DS individuals, their N partners were isolated after pairing, and the number of pigmented offspring in the N individuals' progeny was counted. The male fertility status of the G1 individual was evaluated using the same protocol. **B:** Individual G1 had a pigmented N father and a DS alb-1 mother. In this case, since the N individual was pigmented, it was not possible to assess the male fertility of the DS individual from this cross. Therefore, we provided a second partner to the mother, after she had laid eggs, and this second partner was alb-2 (in light pink) so that outcrossed offspring would be recognised (alb-1 and alb-2 are complementary lines and produce pigmented babies upon cross-fertilization).

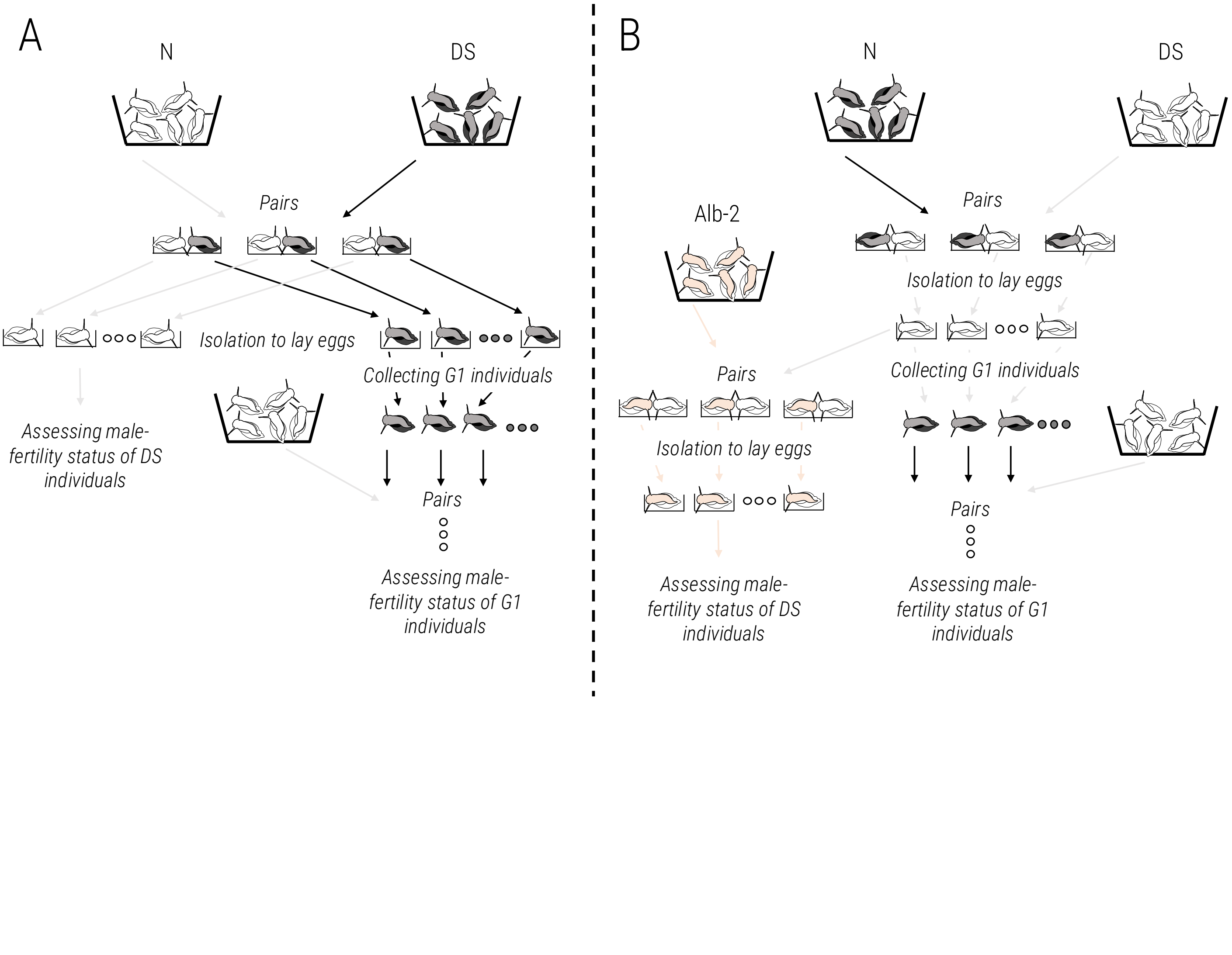

### S4: Additional plots

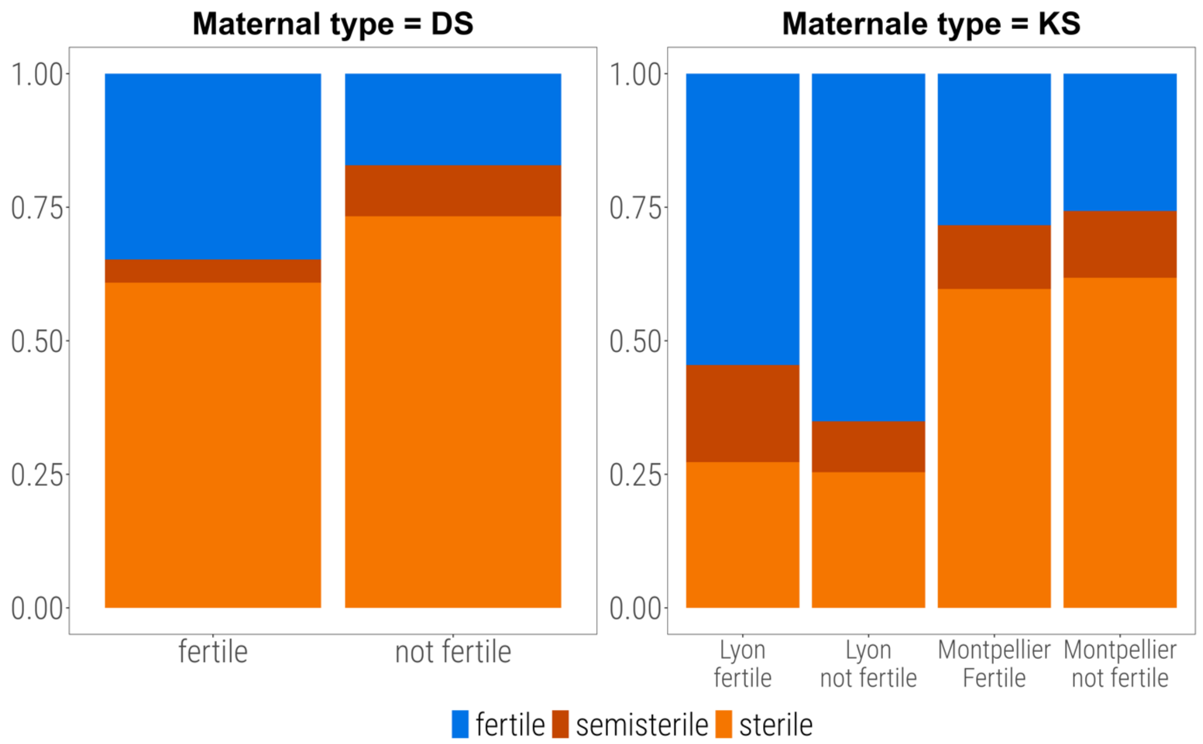


**Figure S4A: Effect of maternal phenotype in the inbred lines dataset.** Only the DS and KS categories are represented because there are not enough male-sterile mothers in the KF category. The number of male-fertile mothers is also not high in the DS category (N=23), so percentages should be taken with caution. The KS dataset has been split by paternal origin because this effect is significant which is not the case in the DS dataset (see main text).

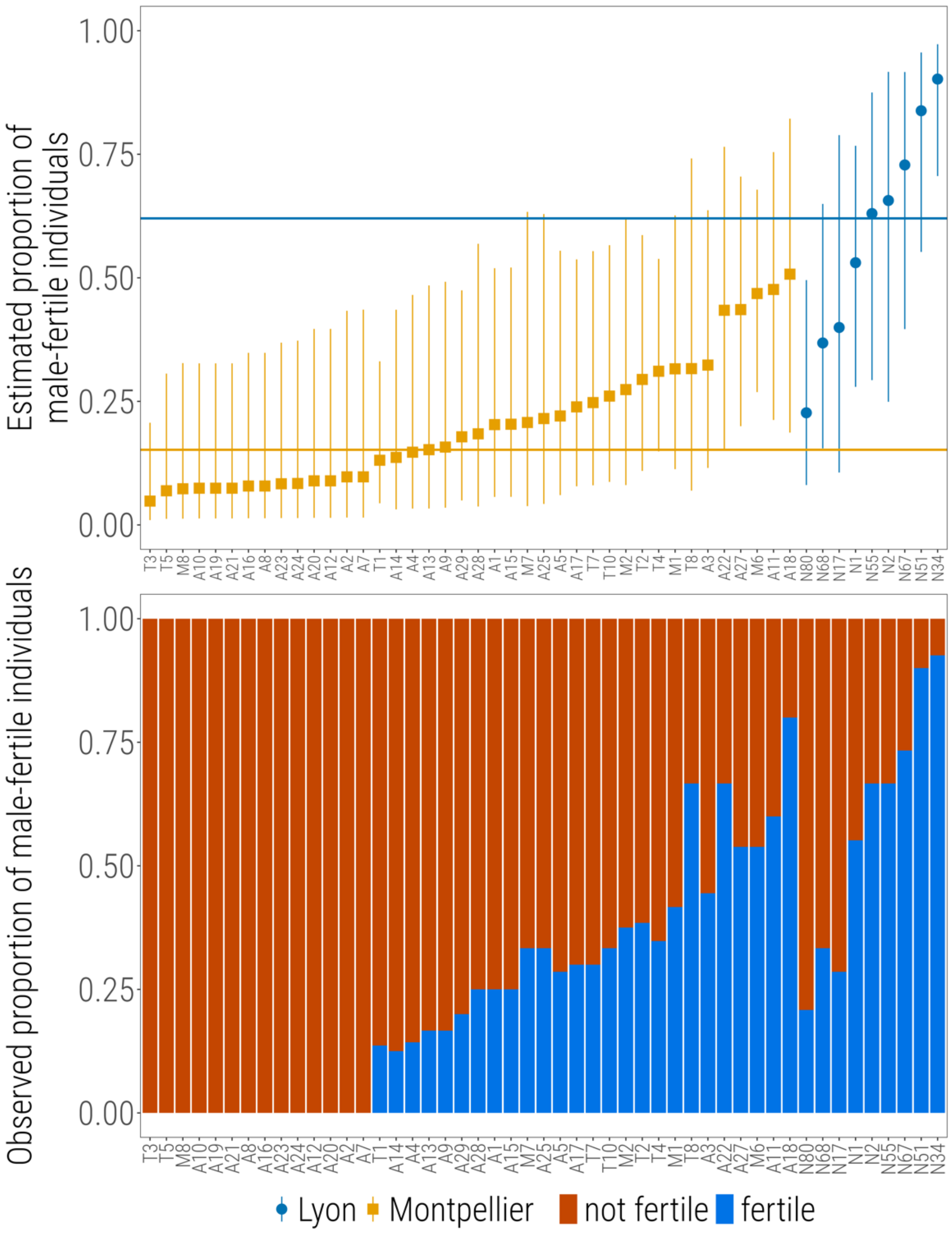


**Figure S4B: Model estimates (top) versus raw proportions (down) of male-sterile individuals for each inbred line crossed with KS mothers.** Inbred lines are arranged in the same order in the two graphs. The figure shows that the continuous distribution of model estimates (BLUPs) is not an artefact of the normality assumption made in the GLMMs, as it is also observed in the raw proportions.

### S5: Heritabilities of male-sterility penetrance under the threshold model

We computed the heritabilities of penetrance of male-sterility under the threshold quantitative-genetic model, separately for each mitotype and population, by re-running the GLMM binomial models on the inbred lines dataset, using the probit link instead of the logit link, as recommended by de Villemereuil et al. 2016. We used twice the estimate of the paternal line variance as our estimate of additive genetic variance in the probit scale, and then obtained the heritability in the natural scale as in de Villemereuil et al. 2016, eq. 24. The complete results are below:

| Data | Random effects | | | | |
|---|---|---|---|---|---|
| | Paternal line variance (probit scale) | Mother variance (probit scale) | % of male-fertiles | $h^2$ (probit scale) | $h^2$ (natural scale) |
| KS, Montpellier | 0.4649 | 0.1773 | 0.25 | 0.539 | 0.291 |
| KS, Lyon | 0.6164 | 0.3278 | 0.5915 | 0.634 | 0.396 |
| DS, Montpellier | 0.2272 | 0.000 | 0.1706 | 0.370 | 0.168 |
| DS, Lyon | 0.2558 | 0.0661 | 0.2462 | 0.387 | 0.207 |
